# Rearing medium dictates variability across replicates in untreated and arsenic challenged zebrafish larvae

**DOI:** 10.1101/2020.08.23.263202

**Authors:** Anjana Ramdas Nair, Patrice Delaney, Shashi Ranjan, Nouf Khan, Catherine Palmer, Kirsten C. Sadler

**Author notes:** To whom correspondence should be addressed: Kirsten C. Sadler, New York University Abu Dhabi, P.O. Box 129188, Abu Dhabi, United Arab Emirates. These authors contributed equally to this work (Anjana Ramdas Nair, Patrice Delaney).

## Abstract

Reproducibility and consistency are hallmarks of scientific integrity. Biological systems are inherently noisy, posing a challenge to reproducibility. This is particularly relevant to the field of environmental toxicology, where many unaccounted experimental parameters can have a marked influence on the biological response to exposure. Here, we extend the use of zebrafish as a robust toxicological model for studying the effects of inorganic arsenic (iAs) on liver biology. We observed that iAs toxicity in this system is not influenced by important parameters including genetic background, rearing container material or rearing volume but the dose response to iAs is influenced by the rearing medium. We compared mortality as a measure of iAs toxicity to embryos cultured in two standard rearing media: egg water made from dehydrated ocean salts dissolved in water and a defined embryo medium which is a pH adjusted, buffered salt solution. Larvae reared in egg water were more susceptible to iAs compared to those reared in embryo medium. This effect was independent of the pH differences between these solutions. These culture conditions did not cause any difference in the global hepatic transcriptome of control zebrafish. Further, no difference in the expression of genes involved in the unfolded protein response (UPR) in larvae exposed to iAs treatment or in a stress independent system to activate UPR genes by transgenic overexpression of activating transcription factor 6 (nAtf6) in hepatocytes was observed. However, the clutch-to-clutch variation in gene expression was significantly greater in larvae reared in egg water compared to those in embryo medium. These data demonstrate that egg water affects reproducibility across replicates in terms of gene expression and exacerbates iAs mediated toxic response. This highlights the importance of rigorous evaluation of experimental conditions to assure reproducibility.

## INTRODUCTION

The reproducibility of scientific findings are the foundation of societal confidence in scientific research. Intensifying efforts and extensive discussions on research reproducibility – or the lack thereof – have characterized the scientific discourse in recent years [1]. One striking report found that over 70% of biological studies could not be reproduced by other research teams and nearly 60% of experiments failed to be reproduced within the same laboratory [1]. Scientific journals, societies and funding agencies are focusing increased effort and resources on identifying the source of variability in science [1, 2]. Many of the recommendations for ensuring rigor and reproducibility focus on improving the detail in the materials and methods that accompany research articles. While much of the variability in experimental outcomes, even within laboratories, can be attributed to the skill and experience of the investigator carrying out the work, experimental parameters that are not controlled for can also have profound effects.

Biological systems are complex, chaotic systems and thus many factors can skew observed results, making the data difficult to interpret and reproduce. As researchers aim to delineate gene expression profiles and their impact on clinically relevant outcomes, there is a community-wide acknowledgement that results can be attributed to unexamined, poorly characterized experimental variables. This is particularly worrisome for toxicological studies, as experimental parameters hold the potential to modify biological responses making it unclear which pathways the toxicant affects. Thus, it is crucial that researchers not only report detailed protocols, but also evaluate the contribution of experimental parameters to the overall discovery.

Model organisms are important tools that not only allow researchers to determine the reproducibility of their findings but also serve to characterize the effects of experimental parameters. Further, *in vivo* models are essential for the translation of toxicological induced clinical manifestations. Although other vertebrate models are more closely related to humans, zebrafish offer many advantages in toxicological studies. First, the rapid development of zebrafish embryos facilitates development of new genetic lines with high statistical precision and assessment of reproducibility compared to other vertebrate models [3]. Second, the community based and comprehensive functional genomic approaches, the widespread use of forward and reverse genetic screens and the excessive characterization of the response to environmental toxicants provides a deep understanding of how vertebrates respond to genetic and chemical perturbations [4]. Third, many organs, including the liver, are fully formed by 120 hours post fertilization (hpf) [5]. The rapid developmental time course coupled with the transparent nature of the larvae allows for quality images of organs from live larvae. Since toxicants can be directly administered to the rearing water, this allows for uniform exposure across larvae which can limit the variability that occurs in other vertebrate models whereby toxicants are administered through food or water, and therefore subject to variable intake. Finally, many metabolic pathways in the liver and other organs are molecularly conserved, making zebrafish a relevant model for toxicant-induced gene expression changes and associated pathologies in humans [6]. These make zebrafish a powerful model for toxicological research and, in particular, the discovery of pathways that mediate liver pathology [7]. Regardless, this model is also susceptible to issues of reproducibility.

Although the power of zebrafish as a model organism is well established, there is a lack of standardization in rearing protocols. Two solutions commonly used in rearing zebrafish larvae during the first five days post fertilization have dramatically different compositions, and the pH and sterilization protocols are highly variable. The first solution, referred to as “egg water,” is a mixture of reverse osmosis (RO) water and dehydrated ocean salts without any additional buffer or supplementary ions and it is not pH adjusted. The second solution, “embryo medium,” is prepared in accordance to the standard Zebrafish Information Network protocol [8] and includes a buffer, calcium and magnesium salts and is adjusted to pH 7.2-7.4. Variability in embryo culture parameters can reduce reproducibility across labs, thereby confounding results. More basic characterization of experimental parameters can help strengthen findings in zebrafish toxicological results.

Our lab has developed a protocol for using of zebrafish as a model for developmental exposure of inorganic arsenic (iAs) [9]. iAs exposure is a serious public health issue, as hundreds of millions of people are at risk of exposure to unsafe levels of iAs through water and food [10]. In many cases, contamination may go unnoticed, posing an insidious risk. Strong data from epidemiological and animal studies have shown that exposure to high levels of arsenic are associated with diabetes, liver disease, cardiovascular disease, cancer, and premature death, and acute to high levels are lethal [11-13]. Hepatocytes are among the most severely affected cells, as the liver is the site where majority of iAs is metabolized [14-23]. Indeed, we found that one stress response pathway, the unfolded protein response (UPR), was activated after chronic (6-120 hpf) iAs exposure in the liver of zebrafish larvae [9].

Here, we expand on our findings and extend the use of zebrafish larvae as an *in vivo* model for iAs toxicological studies by evaluating multiple factors in rearing protocols that may confound iAs induced morality and gene expression in larval livers. We find that many basic parameters, such as genetic background, individual mating pair, rearing container material and volume of water at a fixed density of fish, have no effect on iAs-induced mortality. However, the rearing solution consistently skews the lethal dose of iAs in which fifty percent of the larvae die (LC_50_) such that egg water reared larvae are more susceptible to iAs challenge. Notably, although no significant differences in mean gene expression changes were detected in larval livers between rearing media, egg water reared samples displayed a higher variance across replicates in contrast to embryo medium. This highlights how attention to detail can make a significant difference in the ability to generate reproducible results, and investigators who fail to adhere to this principle can run the risk of failure to reproduce results, even those generated by members of the same laboratory. These findings demonstrate that the components of rearing solution in zebrafish toxicological studies can influence the responses to iAs and can affect reproducibility across replicates in terms of gene expression.

## RESULTS

### iAs toxicity in zebrafish is independent of genetic background, rearing container, and volume of culture media

To expand on previous work establishing zebrafish larvae as an effective model system to study the cellular basis of iAs hepatoxicity [9, 24-31], we examined the genetic strain and culture condition as potential confounding factors that might influence iAs susceptibility in this system. We used mortality as an indicator of systemic iAs toxicity. In previous studies [9] we used a chronic treatment model whereby embryos were exposed to iAs throughout development (6-120 hpf). We used this exposure scheme to test experimental parameters that could potentially influence susceptibility to iAs concentrations ranging from 0.4-2 mM from 6-120 hpf by measuring mortality. We use the inbred strains TAB5, TAB14 and the AB strain (called ABNYU in our facility) and found that embryos from these different genetic backgrounds did not differ in their susceptibility to iAs when we assessed the offspring of group matings (Figure 1A) nor when we compared offspring from mating single pairs of ABNYU adults (Figure 1B). This suggests that in zebrafish, iAs toxicity is consistent across these genetic backgrounds. Additionally, plastics can interact with some toxicants, but we found no difference in the response of embryos cultured in plastic or glass containers to iAs (Figure 1C). Since gas distribution can fluctuate based on the surface area of the media and animal density, we tested whether the culture affected susceptibility by adjusting the volume in a multi-well plate using a fixed density of larvae (2 larvae per mL rearing medium), and found no difference in mortality across these conditions (Figure 1D). Therefore, the parameters of genetic background, rearing container material and volume do not affect iAs-induced toxicity in zebrafish larvae.

**Figure 1.**
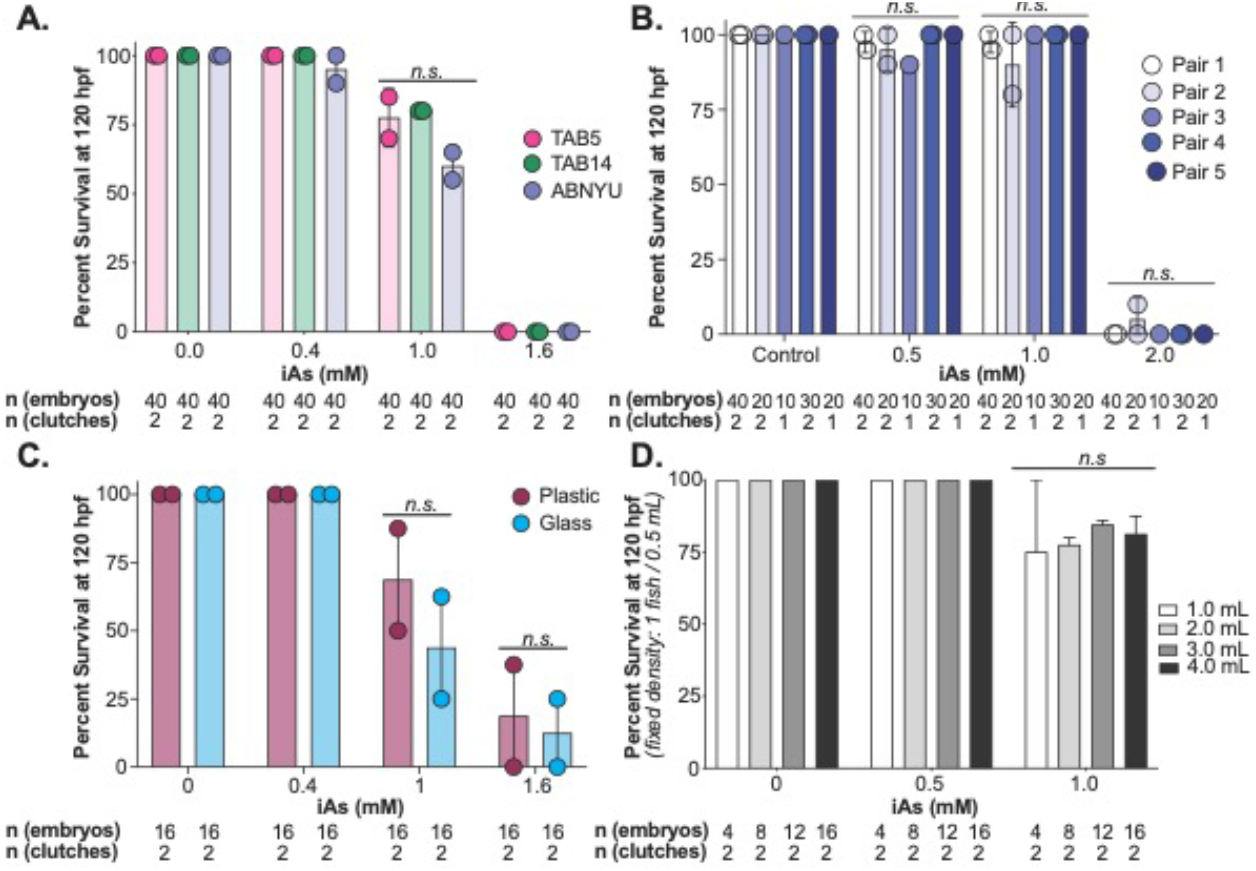
iAs induced toxicity in zebrafish is independent of genetic background, plastic interactions, and volume. **A.** Survival of zebrafish from TAB5 (pink), TAB14 (green) and ABNYU (blue) genetic background at 120 hpf after treatment from 6-120 hpf with 0, 0.4, 1.0 and 1.6 mM of iAs. Dots represent individual clutch values and bars indicate the mean between multiple clutches. *n.s.* indicates no significance, 2-way ANOVA. **B.** Survival analysis of single pairs of ABNYU zebrafish pairs treated with iAs (0 – 2.0 mM). No single pair exhibited resistance or sensitivity to iAs (n.s. = no significance per 2-way ANOVA). **C.** Survival analysis of zebrafish reared in plastic (maroon) and glass (light blue) containers. *n.s.* indicates no significance by 2-way ANOVA. **D.** Survival analysis of zebrafish reared in a fixed density (1 fish/0.5 mL) in volumes ranging from 1-4 mL and exposure to iAs (6-120 hpf, 0-1 mM). *n.s.* indicates no significance, 2-way ANOVA.

### Embryo rearing solution alters the lethal concentration of acute and chronic iAs challenge

We next examined the impact of rearing solution on iAs challenge in zebrafish larvae (Table 1). We hypothesized that the addition of buffer in embryo medium would maintain a stable pH overtime, whereas the pH of unbuffered egg water would fluctuate. Freshly prepared egg water was not only initially more acidic, but gradually increased pH overtime (Figure 2A).

**Table 1:** Egg water and embryo medium constituents.

| Table 1: Egg water and embryo medium constituents |  |  |
| --- | --- | --- |
| Egg Water |  |  |
| Reagent | Quantity | Final Concentration |
| Crystal salt | 0.6 g | 0.06 g/L |
| RO Water | 10 L | Not applicable |
| pH | 6.8-7.2 |  |
| Embryo Medium |  |  |
| Reagent | Quantity | Final Concentration |
| NaCl | 4 g | 13.7 mM |
| KCl | 0.199 g | 0.54 mM |
| Na<sub>2</sub>HPO<sub>4</sub> | 0.178 g | 0.025 mM |
| CaCl<sub>2</sub> | 1.4 g | 3 mM |
| MgSO<sub>4</sub>·7H<sub>2</sub>O | 2.453 g | 2 mM |
| NaHCO<sub>3</sub> | 3.486 g | 4 mM |
| KH<sub>2</sub>PO<sub>4</sub> | 0.299 g | 0.044 mM |
| Autoclaved RO Water | 5 L | Not applicable |
| pH | 7.2-7.3 |  |

**Figure 2.**
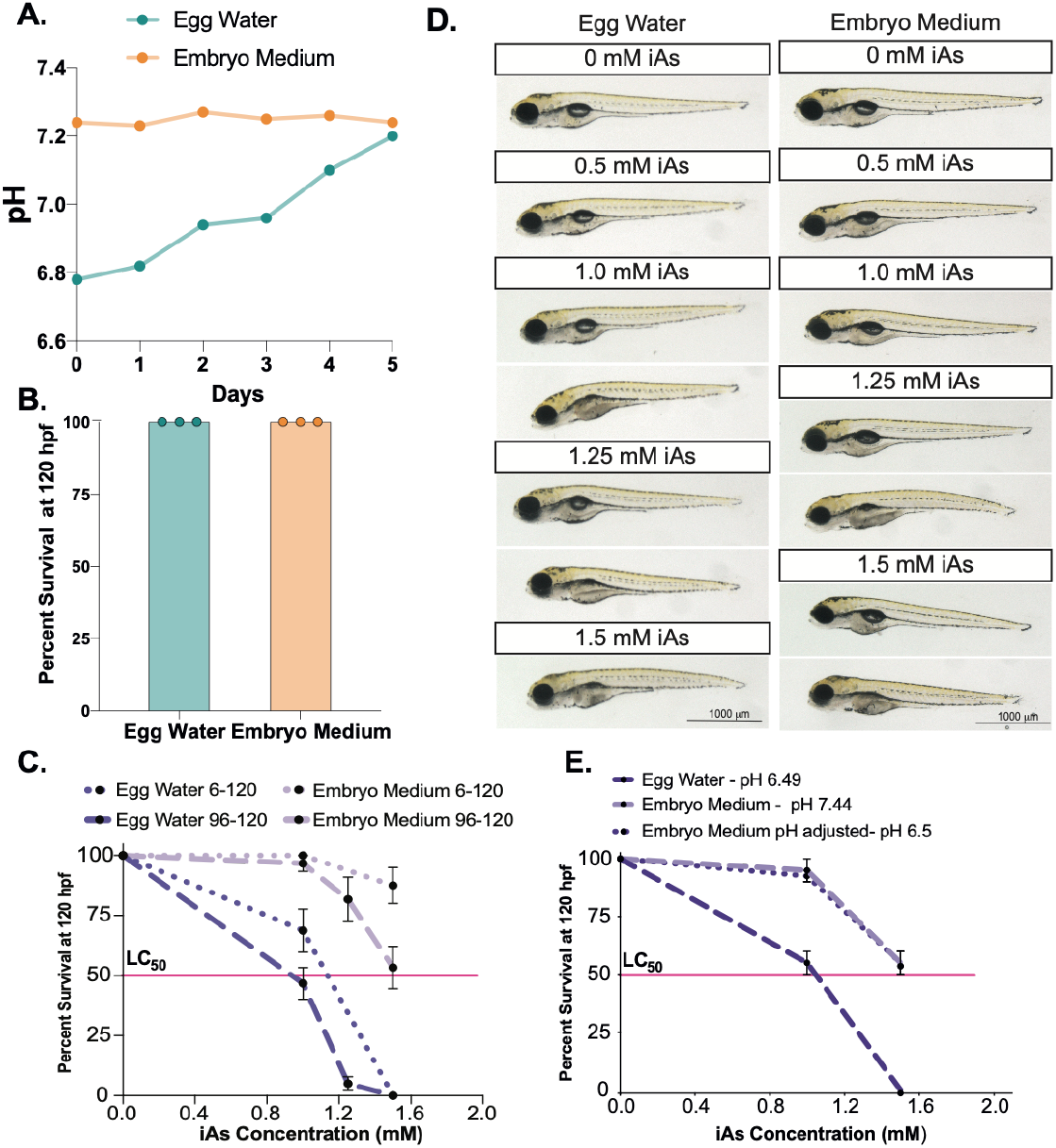
iAs toxicity is dependent on the rearing medium but independent of pH changes. **A.** pH reads of egg water (teal) and embryo medium (orange) monitored daily over the course of 5 days. **B**. Percent survival of zebrafish embryos reared in egg water (teal) or embryo medium (orange) from 6 – 120 hpf. **C**. Surival curve of zebrafish larvae reared in embryo medium (pH 7.2, + Ca and Mg) or egg water (not buffered, mix of ocean salts) exposed to iAs (0-2.0 mM) from 6-120 hpf and 96-120 hpf. Pink line indicates the lethal concentration that causes 50% death (LC_50_). **D.** Representative images of zebrafish at 120 hpf reared in egg water or embryo medium) and exposed to different concentrations of iAs. **E.** Surival curve of zebrafish larvae reared in egg water (pH 6.49), embryo medium (pH 7.4) or embryo medium with pH adjusted (pH 6.49) exposed to iAs (0-1.5 mM) from 96-120 hpf. Pink line indicates the lethal concentration that causes 50% death (LC_50_).

Conversely, fresh embryo medium had a neutral pH that was maintained over a five-day period (Figure 2A). This suggests that embryo medium will produce more consistent results across experimental replicates whereas egg water may cause more variation due to pH changes.

The effects of rearing medium were assessed on the morphology and mortality of zebrafish larvae during both standard rearing conditions and during iAs challenge (Fig. 2B-D). As expected, there were no differences in the survival of fertilized embryos or larvae morphology when reared in egg water or embryo medium (Figure 2B, D). However, we found that the culture medium significantly impacts larval mortality and morphology in response to iAs challenge (Figure 2C-D). During chronic (6-120 hpf) iAs challenge, the concentration at which 50 percent of the larvae die (LC_50_) increases from ∼ 1 mM iAs in egg water to beyond 2 mM iAs in embryo medium (Figure 2C). While the LC_50_ is roughly the same in both egg water and embryo medium during acute iAs challenge (96-120 hpf), 1.5 mM iAs is much better tolerated in embryo medium than in egg water (Figure 2C-D). It is also interesting to note that chronic exposure is better tolerated than the acute exposure, regardless of culture medium.

Given that iAs speciation is sensitive to pH, [32] we tested whether the observed difference in rearing solution can be attributed to pH. Egg water, which has a pH of 6.49, is slightly more acidic than embryo medium, which has a pH of 7.44. We tested the effect of pH on iAs toxicity by generating embryo medium with the same pH of egg water through the addition of hydrochloric acid and found that the LC_50_ of iAs was identical in embryo medium at pH 7.44 and at pH 6.5. This suggests that the observed differences in rearing solution are not due to the acidic pH (Figure 2E).

### iAs toxicity in zebrafish embryos is enhanced following liver development in both egg water and embryo medium

The observation that the LC_50_ of iAs is lower for the acute exposure protocol compared to the chronic protocol suggested that the developmental stage of iAs exposure affected outcome. We sought to compare the effects of egg water and embryo medium on iAs toxicity during different developmental stages using “addition/subtraction” experiments (Figure 3A-B) whereby iAs was added or removed sequentially to embryos in egg water or embryo medium across the developmental stages encompassing gastrulation (6 hpf) to full larval maturity (120 hpf). Larvae were scored at 120 hpf for mortality and morphological defects. We selected 2 mM iAs as the standard concentration in this series of experiments since this exceeds the LC_50_ for the acute exposure protocol in both egg water and embryo medium (Figure 2C), thereby allowing for binary scores of survival and ease of comparison between rearing media.

**Figure 3.**
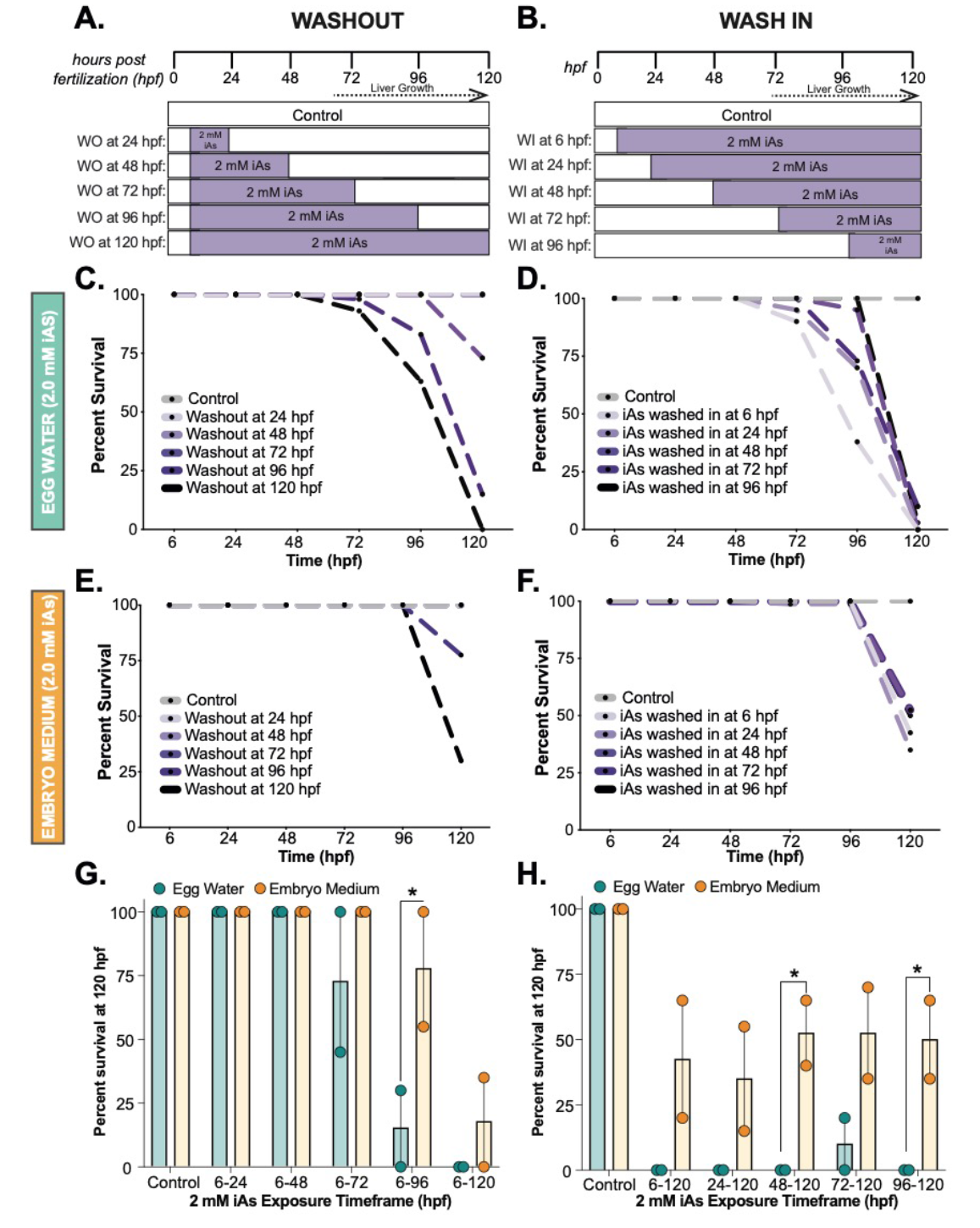
Larval recovery from iAs treatment is decreased after liver bud formation in both egg water and embryo medium. **A.** iAs washout (WO) treatment strategy, whereby all embryos are treated with 2 mM iAs at 6 hpf and iAs is washed out at 24-hour intervals. Animals were monitored until 120 hpf. **B.** iAs wash in (WI) treatment strategy, whereby 2 mM iAs is added at 24-hour intervals and retained in the culture media until 120 hpf. **C.** Survival of zebrafish reared in egg water treated with 2 mM iAs as outlined in A. Fish were scored daily for mortality (n= 40, 2 clutches). **D.** Survival of zebrafish reared in egg water treated with iAs as outlined in B. Fish were scored daily for mortality (n= 40, 2 clutches). **E.** Survival of zebrafish reared in embryo medium treated with iAs as outlined in A. Fish were scored daily for mortality (n= 40, 2 clutches). **F.** Survival of zebrafish reared in embryo medium treated with iAs as outlined in B. Fish were scored daily for mortality (n= 40, 2 clutches). **G.** Bar plots of mean percent survival of WO treatment (A) in egg water (teal) versus embryo medium (orange). Each point represents a single clutch (n= 20 fish). Error bars represent the standard error mean (SEM). * indicates p-value < 0.05, 2way ANOVA. **H.** Bar plots of mean percent survival of WI treatment (B) in egg water (teal) versus embryo medium (orange). Each point represents a single clutch (n= 20 fish). Error bars represent the standard error mean (SEM). * indicates p-value < 0.05, 2way ANOVA.

Exposure to toxicants that directly impair key cellular processes or those causing genomic instability typically causes death during acute exposure, as we recently reported with zebrafish embryos exposed to metals configured in trefoil knots [33]. In contrast, most death in response to iAs exposure occurred between 96-120 hpf, regardless of the time of administration or removal of iAs or rearing medium (Figure 3C-F). This shows that iAs toxicity in zebrafish larvae is not a consequence of a developmental abnormality or cumulative effects. Instead, we concluded that some feature of mature larvae renders them hypersensitive to iAs. This is supported by the finding that 2 mM iAs exposure caused death and morphological abnormalities even if iAs was removed after 96 hpf (Figure 3D-F).

A simple interpretation of the subtraction experiments (Figure 3A, C and E) is that toxicity is directly related to duration of the exposure. To test this, we carried out addition experiments whereby iAs exposure started at 6, 24, 48, 72 or 96 hours and maintained in the water until 120 hpf (Figure 3B). There was death of nearly all larvae in all treatment groups by 120 hpf, even if the exposure duration was as short as 24 hours in either medium (Figure 1D and F). This shows that a concentration of iAs that is lethal in older larvae is well tolerated by early embryos, and demonstrate that the window of iAs susceptibility occurs after 72 hpf in both egg water and embryo medium. Thus, iAs-induced mortality observed in embryos chronically exposed to iAs is not likely to be caused by the developmental effects of exposure, but instead due to the effects of iAs on tissues or cell types present at later developmental stages. This phenomenon is consistent in both rearing media, though occurs slightly earlier in egg water versus embryo medium. As expected, larvae reared in egg water showed higher susceptibility to 2 mM iAs challenge (Figure 3G-H), though did not show death until after liver bud development (Figure 3C-D) [5].

These data support the hypothesis that iAs toxicity is enhanced when the larvae develop hepatic function, and this finding is consistent across egg water and embryo medium. Moreover, these experiments demonstrate that iAs toxicity during chronic exposure is largely due to effects sustained during the latter part of the exposure. These data are consistent with findings in humans and other animal models showing that iAs is a hepatotoxicant [14, 22, 34-36] and suggest that hepatic toxicity is responsible for lethality in this system, regardless if larvae are reared in egg water or embryo medium.

### Larvae reared in egg water have higher variance in gene expression in the liver

As previously described, the rearing medium either confers iAs susceptibility in egg water or iAs resistance in embryo medium. One hypothesis is that egg water introduces a component that increases basal levels of stress response pathways, such as the UPR, thereby overloading stress response capacity upon secondary challenge. Thus, we performed qPCR to test key UPR genes, *hspa5, hyou1* and *atf6*, on livers from larvae reared in egg water or embryo medium. Surprisingly, we found that the egg water had a higher variance and mean; albeit not significant in comparison to embryo medium (Figure 4A). This suggests that egg water does not significantly alter basal UPR expression, but instead confers higher variability in biological response across replicates.

**Figure 4.**
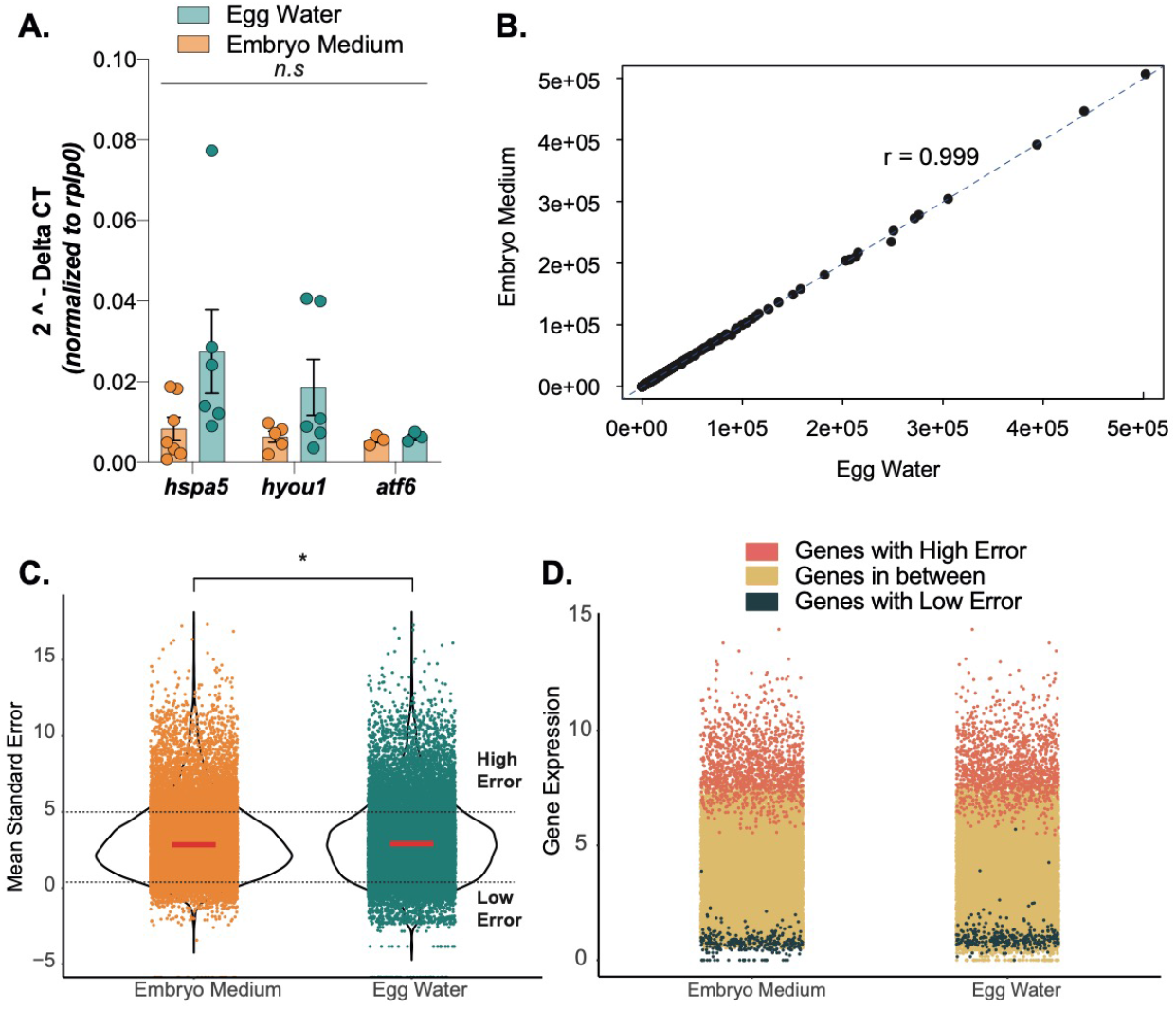
Rearing medium does not affect gene expression but impacts the mean standard error across the samples. **A.** Gene expression of UPR genes *hspa5, hyou1* and *atf6* derived from qPCR analysis on pools of 5 larval zebrafish livers reared in egg water or embryo medium. *n.s* indicates no significance. **B.** Linear correlation plot depicting mean reads across control 120 hpf larvae reared in egg water versus embryo medium. r is the Pearson correlation coefficient. **C.** Jitter plot showing the distribution of mean standard error in log scale of genes expressed in control 120 hpf larvae reared in egg water or embryo medium. The red line indicates the median. **D.** Jitter plot showing the gene expression (log(fpkm+1)) of genes segregated based on standard error in the sample. High error (log(standard error mean)>=5) in coral, Low error (log(standard error mean)<=0.5) in dark green and in between high and low errors in mustard yellow.

To determine if this variability was systematic across all genes, RNAseq from two independently generated control datasets from 5 replicates of pools of livers dissected from 120 hpf larvae that were reared in egg water or embryo medium were analyzed. These samples were collected by the same investigator, and processed in the same genomics facility on different days. These datasets were batch corrected and normalized to generate the Fragments Per Kilobase of transcript per Million mapped reads (FPKM). We found the mean gene expression in these datasets as shown by the Pearson correlation coefficient of 0.999 (Figure 4B). We next asked if there was a difference between replicates in the samples cultured in egg water or embryo medium by determining the standard error of the mean for all genes across biological replicates. This showed that there were more genes with high error in samples cultured in egg water than embryo medium (Figure 4C). As expected, the genes with high error (log(mean standard error) >= 5) among replicates were highly expressed, while genes with low error (log(mean standard error) <= 0.5) across replicates had very low expression (Figure 4D), implicating that the observed range in variance is not due to technical artifacts.

Taken together, these data indicate that rearing medium does not affect overall gene expression, but culturing larvae in egg water increases the variance between samples, conferring higher error values across replicates. This can affect the statistical analysis of samples reared in egg water. Therefore, using embryo medium to rear larvae would lead to more reproducible gene expression values across biological replicates. Although we controlled for investigator mediated differences in this dataset, the individual differences between lab personnel preparing samples can also be a source of variability in other datasets.

### Rearing medium sensitizes livers to iAs exposure-associated UPR response

To determine if the rearing media affected the transcriptional response in the liver to iAs exposure, we focused on the UPR, which we previously demonstrated is altered in zebrafish treated with iAs and other chemical and genetic perturbations that cause endoplasmic reticulum (ER) stress [9, 37-39]. qPCR analysis of liver samples of larvae exposed to iAs (1.0 and 1.5 mM) reared in egg water (pH 6.49), embryo medium (pH 7.4) or embryo medium that has been pH adjusted to match egg water (embryo medium, pH 6.5) showed that the induction of UPR genes was higher in response to 1.0 mM iAs exposure of larvae reared in egg water compared to embryo medium (Figure 5A). In concordance with this finding, 1.5 mM iAs-challenged larvae reared in egg water could not be evaluated for gene expression as all larvae in this cohort died, in contrast, to over 50% survival of larvae reared in embryo medium treated with the same iAs concentration (Figure 2E). Adjusting the pH of embryo medium had no effect on UPR gene expression in basal conditions or in response to iAs (Figure 5A). This suggests that the pH differences between rearing media do not account for the transcriptomic response to ER stress in the liver caused by iAs challenge.

**Figure 5.**
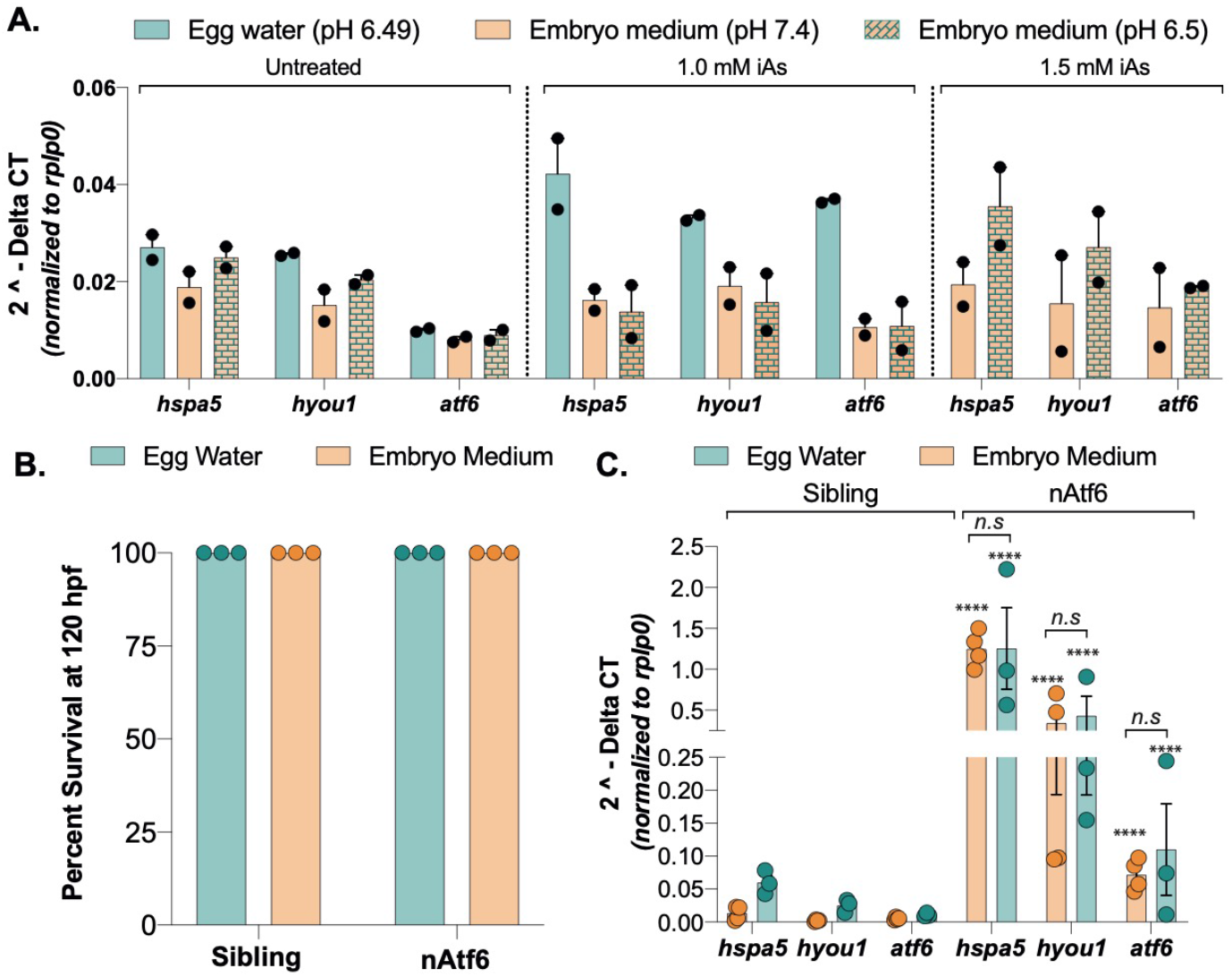
UPR gene expression is upregulated at lower concentrations of iAs when fish are reared in egg water compared to embryo medium and nAtf6 A. Gene expression of UPR genes *hspa5, hyou1* and *atf6* derived from qPCR analysis depicted as 2^-dCt^ values normalized to *rplp0* on pools of 5 larval zebrafish livers reared in egg water (pH 6.49), embryo medium (pH 7.4) or embryo medium with pH adjusted (pH 6.49) that were exposed to 0, 1.0 and 1.5 mM iAs from 96-120 hpf. Each dot corresponds to a clutch. **B.** Bar plots of mean percent survival at 120 hpf of zebrafish larvae that are either *Tg(fabp10a:nAtf6-mCherry)* or sibling reared in egg water or embryo medium. (n = 3 clutches, 60 fish). **C.** Gene expression of UPR genes *hspa5, hyou1* and *atf6* derived from qPCR analysis depicted as 2^-dCt^ values normalized to *rplp0* on pools of livers expressing *Tg(fabp10a:nAtf6-mCherry)* or siblings. *n.s* indicates no significance.

We extended this analysis to a model we previously described [38] in which UPR gene regulation is altered in stress-independent conditions by overexpression of Atf6, a key transcriptional mediator of the UPR, in zebrafish hepatocytes (*Tg(fabp10a:nAtf6-mCherry*). We reared these larvae in egg water and embryo medium and found that there was no effect on survival (Fig. 5B) or phenotype (not shown). Moreover, qPCR analysis on liver samples of siblings and nAtf6 overexpressing larvae showed that, as predicted, aft6 expression was elevated due to overexpression of the transgene, and also there was significant induction of the Atf6 targets *hspa5* and *hyou1*. However, the rearing media had no significant impact on the expression of UPR genes in this model (Fig. 5C). We did observe, however, a greater variation in the range of gene expression in the liver of larvae reared in egg water compared to those reared in embryo medium, reflected by a higher standard error. This indicates the response to ER stress or stress signaling pathways is not significantly altered by the embryo rearing medium.

Further,while the UPR gene expression was preserved, there was higher variability within the replicates as observed in the RNAseq datasets (Figure 4C-D, egg water). Therefore, rearing medium has no significant effect on UPR gene expression in stress free UPR activated models.

Collectively, these results indicate that raising zebrafish larvae in egg water confers more variability in basal gene expression in the liver compared to embryo medium. Further, iAs-treated larvae are more susceptible to iAs at lower doses and tend to have higher UPR gene expression in livers when reared in egg water instead of embryo medium. These differences are specific to toxicant-induced stress and are not observed in stress-independent conditions where Atf6 is activated.

## DISCUSSION

In summary, we optimized a treatment scheme using zebrafish where multiple confounding factors are controlled, providing a robust system for experiments addressing mechanisms of iAs toxicity in the liver. We find that iAs mortality is independent of many basic experimental parameters, but significantly altered by rearing solution. Larvae reared in egg water have a lower LC_50_ for iAs provided as both acute and chronic exposures compared to siblings reared in embryo medium. Despite these differences, we find that embryos and larvae are more susceptible to iAs exposure at late developmental stages, coinciding with the time when hepatocytes differentiate, independent of the solution they are reared in. This suggests that the presence of a mature liver is a factor in conferring iAs toxicity. Gene expression analysis in larval livers revealed that although overall gene expression is the same in larvae raised in the two media examined here, unbuffered egg water introduces more variability. Further, egg water confers greater sensitivity and variability in UPR induction in the liver during iAs challenge, but not during stress-independent UPR activated larval livers. This suggests that the observed rearing medium differences are not due to basal pathway differences in the liver.

We were surprised to find that the different standard rearing media routinely used for zebrafish embryos altered the susceptibility to iAs, as we have not found similar effects in response to other chemicals (not shown). Despite the difference in the LC_50_ for iAs conferred by rearing medium, the response to iAs appears to cause the same phenotype in exposed larvae. Although egg water does not seem to cause differentially expressed genes compared to embryo medium, it is possible that egg water, which is derived from evaporated seawater, could contain trace elements or other toxicants that could sensitize to iAs toxicity. While egg water is simple to make, since the components are not defined, it is difficult to identify a potential source of the variability in this solution.

Another possibility is that embryo medium provides an essential component, such as Ca^2+^ or Mg^2+^, which may provide an important supplement for hepatocellular function. Indeed, hypomagnesemia results in many disorders, including fatigue, seizures and muscle cramps [40]. Zebrafish primarily obtain their intake of magnesium from absorption through the water [40]. Additionally, other studies have shown that zebrafish larvae body length are positively correlated with bioavailability of calcium and magnesium in the rearing medium [41]. These data suggest the addition of mineral supplements in embryo medium could increase biological fitness and therefore may underlie the observed iAs resistance. In rodent models, a variety of nutritional protocols exist to diminish unwanted secondary affects from small molecular inclusions in their food [42]. Since zebrafish larvae are sustained by the nutrients in their yolk during the first 120 hpf, externally provided nutrients during early development has not been considered to be a source of variability. Moreover, zebrafish embryos are considered hearty, and are amenable to rearing under a wide range of water quality [43] and thus little attention has been paid to the nutrition of the early embryo or the components of the defined media that are important for development, growth and response to stimuli. However, other factors that influence nutritional supplementation, such as the availability of minerals in the rearing solution, holds the potential to influence toxicological responses.

An alternative hypothesis is that embryo medium may cause iAs resistance through chelation of iAs, reducing the exposure. Interestingly, calcium and magnesium are routinely used to immobilize arsenates in water treatment plants [44]. While no precipitates were observed in embryo medium during our experiment, this raises the possibility that iAs interactions with calcium and magnesium ions could reduce the effective dose of iAs. More biochemical studies are necessary to evaluate the interactions between iAs and rearing medium ion interactions. Defining the precise mechanism underlying the different effects of these media on iAs toxicity requires further investigation.

Finally, it is possible that egg water could increase iAs susceptibility because of its debilitating effect on other organs. Studies have shown the importance of pathways involved in magnesium uptake in renal maintenance [40]. Another important mineral, calcium is required in bone formation, cardiac and kidney function [42]. In our study, while we did observe an effect in mortality at lower concentrations of iAs in fish reared in egg water, this could not be attributed to compromised liver function (data not shown). Therefore, it is possible other organs, such as the kidney or the heart, could confer the sensitivity to secondary iAs challenge in egg water.

Our work focuses on the liver, since hepatocytes are the site of xenobiotic metabolism and detoxification, making it particularly vulnerable to iAs toxicity. Indeed, our study demonstrates that susceptibility to iAs toxicity in zebrafish larvae is correlated with the formation of the liver, suggesting that the liver is the main contributor to the systemic iAs toxicity. These data reflect the clinical manifestations in humans, as populations exposed to iAs are at increased risk for liver disease and cancer [12, 14, 17, 20-23, 34, 45, 46]. Although further investigation is necessary to reveal how iAs metabolism in the liver can potentiate systematic toxicity, this study provides preliminary evidence that zebrafish are a powerful model for iAs induced liver disease. Further, we provide a robust protocol for future investigations in zebrafish larvae in iAs toxicity.

In summary, this study demonstrates that attention to detail in biomedical science matters, and that reproducibility can be influenced by factors that are difficult to identify. While there is a crisis of reproducibility in many branches of science [1], environmental toxicology is perhaps one of the most susceptible, as many unrecognized factors can synergize, interact to alter the responses to toxicants. While the approaches considered here are extensive, they are by no means exhaustive, and further work is warranted to identify potential factors that influence rigor and reproducibility in experimental systems. However, regardless of experimental parameters, the experience in our own research group is that variability of experimental outcomes and failure to reproduce published results is directly related to the technical expertise of the investigator: no matter how stringent the rearing conditions, the ability of the researcher to carry out careful, well controlled and, at times, labor intensive, experimental protocols is an important factor underling the ability to generate reproducible results.

## MATERIALS & METHODS

### Zebrafish husbandry and embryo rearing

All procedures were approved by and performed in accordance with the New York University Abu Dhabi Institutional Animal Care and Use Committee (IACUC) guidelines. Adult wild type (WT; ABNYU, TAB5, and BNY14) and transgenic lines *(Tg(fabp10a:nls-mCherry)* and *Tg(fabp10a:Gc-EGFP))* were maintained on a 14:10 light:dark cycle at 28°C. All experiments were conducted in 6-well plates (Corning, USA) with 20 embryos in 10 mL rearing medium. Embryos were collected from group matings or single mating pairs, as indicated, within 2 hours of spawning and were reared at 28 °C according to standard conditions.

All experiments were carried out with embryos and larvae reared in embryo medium, unless indicated. The working solution of embryo medium and egg water was prepared fresh weekly as detailed in Table 1.

### Embryo exposure

Embryos were exposed to sodium meta-arsenite (Sigma-Aldrich, USA; heretofore referred to as iAs) by diluting 0.05 M stock solution to final concentrations ranging from 0.4 mM - 2 mM in egg water or embryo medium, as indicated in the text. For addition experiments, 2.0 mM of iAs was added to embryos at 6, 24, 48, 72 and 96 hpf. For subtraction experiments, 2.0 mM iAs was added at 6 hpf and removed at 24, 48, 72, 96 and 120 hpf by transferring embryos in 10 mL of fresh embryo medium twice. For all experiments, mortality was scored daily, dead embryos were removed upon identification, and morphology was recorded at 120 hpf as readout of iAs toxicity.

### Image Acquisition

For whole mount imaging of live larvae, embryos were anesthetized with 500 μM tricaine (Ethyl 3-aminobenzoate methanesulfonate; Sigma-Aldrich), mounted in 3% methyl-cellulose on a glass slide and imaged on a Nikon SMZ25 stereomicroscope.

### Gene Expression Analysis

Pools of at least 5 livers were microdissected from 120 hpf zebrafish larvae with transgenic marked livers (*Tg(fabp10a:DsRed;ela:eGFP))* or (*Tg(fabp10a:Caax-eGFP))*. Larvae were anesthetized in tricaine and immobilized in 3% methyl cellulose and the livers were removed using 30-gauge needles. RNA was extracted from livers using TRIzol (Thermo Fisher, 15596026) and precipitated with isopropanol as described [47]. RNA was reverse transcribed with qScript (QuantaBio, 95048-025).

Gene expression was assessed using RNAseq or quantitative reverse transcription PCR (qRT-PCR). qRT-PCR was performed using Maxima Sybr Green/ROX qPCR Master Mix Super Mix (Thermo Fisher, K0221). Samples were run in triplicate on QuantStudio 5 (Thermo Fisher). Target gene expression was normalized to *ribosomal protein large P0* (*rplp0*) using the comparative threshold cycle (ΔCt) method. Expression in treated animals was compared to untreated controls from the same clutch to determine fold change.

RNAseq analysis was done on the untreated samples from two independent datasets where larvae from one sample set were reared in egg water (Gene Expression Omnibus (GEO) accession number GSE151291) and the other was reared in embryo medium (GEO accession number GSE156420). To adjust the batch effect, we adopted a method designed for RNA-seq count data implemented by Combat-seq [48]. The method keeps the negative binomial distribution of RNA-seq reads count and the integer nature of the data. Then corrected gene counts are feed to DESeq2 for differential gene analysis.

### Statistical analysis, rigor and reproducibility

All experiments were repeated on at least 2 clutches of embryos when possible, with all replicates indicated. Reproducibility was assured by carrying phenotype scoring and other key experiments by independent investigators. Data are presented as normalized values and, where appropriate, raw data is provided. Statistical tests were used as appropriate to the specific analysis, including Student’s T-test, ANOVA and Chi Square (Fisher’s exact test) using Graphpad Prism Software. Linear correlation plot and the Jitter plots were plotted using R to analyze the batch corrected data.

## Author Contributions

ARN, PD and KCS conceived the idea and planned the experiments. ARN, PD, SR, CP and NK carried out the experiments and KCS, ARN, PD, SR, CP and NK analyzed the data and prepared the figures and wrote the manuscript.

## Acknowledgements

The authors are grateful to K. Bambino for preliminary data and to members of the Sadler lab, especially Chi Zhang, for input and technical support. Additionally, the authors are indebted to the NYUAD Core Technology Platform Imaging Facility, especially R. Rezgui, M. Sultana, M. Arnoux, and N. Drou, for continued technical support and careful maintenance of the facilities.

## Notes

The authors declare no conflict of interest.

### Competing Interest Statement

The authors have declared no competing interest.

## References

1. Baker, M., 1,500 scientists lift the lid on reproducibility. Nature, 2016. 533(7604).

2. The challenges of replication. eLife, 2017. 6: p. e23693.

3. Meyers, J.R., Zebrafish: Development of a Vertebrate Model Organism. Current Protocols Essential Laboratory Techniques, 2018. 16(1): p. e19.

4. Lawson, Nathan D. and Scot A. Wolfe, Forward and Reverse Genetic Approaches for the Analysis of Vertebrate Development in the Zebrafish. Developmental Cell, 2011. 21(1): p. 48–64.

5. Chu, J. and K.C. Sadler, New school in liver development: lessons from zebrafish. Hepatology, 2009. 50(5): p. 1656–63.

6. Palmgren, M., et al., AS3MT-mediated tolerance to arsenic evolved by multiple independent horizontal gene transfers from bacteria to eukaryotes. PloS one, 2017. 12(4): p. e0175422–e0175422.

7. Goessling, W. and K.C. Sadler, Zebrafish: an important tool for liver disease research. Gastroenterology, 2015. 149(6): p. 1361–77.

8. Westerfield, M., The zebrafish book. A guide for the laboratory use of zebrafish (Danio rerio). The Zebrafish Book. 2000, Univ. of Oregon Press, Eugene: Zebrafish International Resource Center.

9. Bambino, K., et al., Inorganic arsenic causes fatty liver and interacts with ethanol to cause alcoholic liver disease in zebrafish. Disease models & mechanisms, 2018. 11(2): p. dmm031575.

10. Liao, N., et al., A Comprehensive Review of Arsenic Exposure and Risk from Rice and a Risk Assessment among a Cohort of Adolescents in Kunming, China. International journal of environmental research and public health, 2018. 15(10): p. 2191.

11. Smith, A.H., E.O. Lingas, and M. Rahman, Contamination of drinking-water by arsenic in Bangladesh: a public health emergency. Bull World Health Organ, 2000. 78(9): p. 1093–103.

12. Naujokas, M.F., et al., The broad scope of health effects from chronic arsenic exposure: update on a worldwide public health problem. Environ Health Perspect, 2013. 121(3): p. 295–302.

13. Kuo, C.C., et al., The Association of Arsenic Metabolism with Cancer, Cardiovascular Disease, and Diabetes: A Systematic Review of the Epidemiological Evidence. Environ Health Perspect, 2017. 125(8): p. 087001.

14. Santra, A., et al., Hepatic manifestations in chronic arsenic toxicity. Indian J Gastroenterol, 1999. 18(4): p. 152–5.

15. Santra, A., et al., Hepatic damage caused by chronic arsenic toxicity in experimental animals. J Toxicol Clin Toxicol, 2000. 38(4): p. 395–405.

16. Guha Mazumder, D.N., Arsenic and liver disease. J Indian Med Assoc, 2001. 99(6): p. 311, 314-5, 318-20.

17. Chen, Y. and H. Ahsan, Cancer burden from arsenic in drinking water in Bangladesh. Am J Public Health, 2004. 94(5): p. 741–4.

18. Kosanovic, M., et al., Influence of urbanization of the western coast of the United Arab Emirates on trace metal content in muscle and liver of wild Red-spot emperor (Lethrinus lentjan). Food Chem Toxicol, 2007. 45(11): p. 2261–6.

19. Arteel, G.E., et al., Subhepatotoxic exposure to arsenic enhances lipopolysaccharide-induced liver injury in mice. Toxicol Appl Pharmacol, 2008. 226(2): p. 128–39.

20. Liaw, J., et al., Increased childhood liver cancer mortality and arsenic in drinking water in northern Chile. Cancer Epidemiol Biomarkers Prev, 2008. 17(8): p. 1982–7.

21. Liu, J. and M.P. Waalkes, Liver is a target of arsenic carcinogenesis. Toxicol Sci, 2008. 105(1): p. 24–32.

22. Hao, L., et al., Hepatotoxicity from arsenic trioxide for pediatric acute promyelocytic leukemia. J Pediatr Hematol Oncol, 2013. 35(2): p. e67–70.

23. Frediani, J.K., et al., Arsenic exposure and risk of nonalcoholic fatty liver disease (NAFLD) among U.S. adolescents and adults: an association modified by race/ethnicity, NHANES 2005-2014. Environ Health, 2018. 17(1): p. 6.

24. Li, D., et al., Developmental mechanisms of arsenite toxicity in zebrafish (Danio rerio) embryos. Aquat Toxicol, 2009. 91(3): p. 229–37.

25. Hamdi, M., et al., Identification of an S-adenosylmethionine (SAM) dependent arsenic methyltransferase in Danio rerio. Toxicol Appl Pharmacol, 2012. 262(2): p. 185–93.

26. Sarkar, S., et al., Low dose of arsenic trioxide triggers oxidative stress in zebrafish brain: expression of antioxidant genes. Ecotoxicol Environ Saf, 2014. 107: p. 1–8.

27. Ma, Y., et al., Folic acid protects against arsenic-mediated embryo toxicity by up-regulating the expression of Dvr1. Sci Rep, 2015. 5: p. 16093.

28. Fuse, Y., V.T. Nguyen, and M. Kobayashi, Nrf2-dependent protection against acute sodium arsenite toxicity in zebrafish. Toxicol Appl Pharmacol, 2016. 305: p. 136–142.

29. Beaver, L.M., et al., Combinatorial effects of zinc deficiency and arsenic exposure on zebrafish (Danio rerio) development. PLoS One, 2017. 12(8): p. e0183831.

30. Sun, H.J., et al., Environmentally relevant concentrations of arsenite induces developmental toxicity and oxidative responses in the early life stage of zebrafish. Environ Pollut, 2019. 254(Pt A): p. 113022.

31. Valles, S., et al., Exposure to low doses of inorganic arsenic induces transgenerational changes on behavioral and epigenetic markers in zebrafish (Danio rerio). Toxicol Appl Pharmacol, 2020. 396: p. 115002.

32. Smedley, P.L. and D.G. Kinniburgh, A review of the source, behaviour and distribution of arsenic in natural waters. Applied Geochemistry, 2002. 17(5): p. 517–568.

33. Benyettou, F., et al., Potent and selective in vitro and in vivo antiproliferative effects of metal-organic trefoil knots. Chemical Science, 2019. 10(23): p. 5884–5892.

34. Mazumder, D.N., Effect of chronic intake of arsenic-contaminated water on liver. Toxicol Appl Pharmacol, 2005. 206(2): p. 169–75.

35. Tan, M., et al., Chronic subhepatotoxic exposure to arsenic enhances hepatic injury caused by high fat diet in mice. Toxicol Appl Pharmacol, 2011. 257(3): p. 356–64.

36. Nain, S. and J.E. Smits, Pathological, immunological and biochemical markers of subchronic arsenic toxicity in rats. Environ Toxicol, 2012. 27(4): p. 244–54.

37. Vacaru, A.M., et al., Molecularly defined unfolded protein response subclasses have distinct correlations with fatty liver disease in zebrafish. Disease models & mechanisms, 2014. 7(7): p. 823–835.

38. Howarth, D.L., et al., Activating transcription factor 6 is necessary and sufficient for alcoholic fatty liver disease in zebrafish. PLoS genetics, 2014. 10(5): p. e1004335–e1004335.

39. Tsedensodnom, O., et al., Ethanol metabolism and oxidative stress are required for unfolded protein response activation and steatosis in zebrafish with alcoholic liver disease. Dis Model Mech, 2013. 6(5): p. 1213–26.

40. Arjona, F.J., et al., SLC41A1 is essential for magnesium homeostasis in vivo. Pflügers Archiv - European Journal of Physiology, 2019. 471(6): p. 845–860.

41. Chang, C. and J. Zhu, Different Effects of Three Types of Water on Developmental Behaviors, Lipid Metabolism and Antioxidant Capacity of Juvenile Zebrafish. bioRxiv, 2018: p. 373381.

42. Watts, S.A., M. Powell, and L.R. D’Abramo, Fundamental approaches to the study of zebrafish nutrition. ILAR journal, 2012. 53(2): p. 144–160.

43. Spence, R., et al., The behaviour and ecology of the zebrafish, Danio rerio. Biol Rev Camb Philos Soc, 2008. 83(1): p. 13–34.

44. Magalhães, M.C.F., Arsenic. An environmental problem limited by solubility. Pure and Applied Chemistry, 2002. 74(10): p. 1843–1850.

45. Smith, A.H., et al., Cancer risks from arsenic in drinking water. Environ Health Perspect, 1992. 97: p. 259–67.

46. Young, J.L., L. Cai, and J.C. States, Impact of prenatal arsenic exposure on chronic adult diseases. Syst Biol Reprod Med, 2018. 64(6): p. 469–483.

47. Vacaru, A.M., et al., Molecularly defined unfolded protein response subclasses have distinct correlations with fatty liver disease in zebrafish. Dis Model Mech, 2014. 7(7): p. 823–35.

48. Zhang, Y., G. Parmigiani, and W.E. Johnson, ComBat-Seq: batch effect adjustment for RNA-Seq count data. bioRxiv, 2020: p. 2020.01.13.904730.

